# Local adaptation to hosts and parasitoids shape *Hamiltonella defensa* genotypes across aphid species

**DOI:** 10.1101/2022.07.01.498469

**Authors:** Taoping Wu, David Monnin, Rene A. R. Lee, Lee M. Henry

## Abstract

Facultative symbionts are common in insects and are known to provide important adaptations that can drive rapid host evolution. Yet we still have a limited understanding of what shapes their distributions, such as why particular symbiont strains are common in some host species yet absent in other. To address this question, we genotyped the defensive symbiont *Hamiltonella defensa* in 26 aphid species that commonly carry this microbe. We found that *Hamiltonella* strains were strongly associated with specific aphid species and that strains found in one host species rarely occurred in others. To explain these associations, we reciprocally transferred the *Hamiltonella* strains of 3 aphid species, *Acyrthosiphon pisum, Macrosiphoniella artemisiae* and *Macrosiphum euphorbiae*, in the other host species, and assessed the impact of *Hamiltonella* strain on: the stability of the symbiosis, aphid fecundity, and parasitoid resistance. We demonstrate that the *Hamiltonella* associations found in nature are locally adapted both to the aphid host itself, and its ecology, in that aphids tend to carry *Hamiltonella* strains that provide strong protection against their dominant parasitoid species. Our results suggest that *Hamiltonella* strains function as a horizontal gene pool that aphids draw from to rapidly adapt to pressures from different natural enemies.

## Introduction

Many insects harbour intracellular bacteria that profoundly influence their biology. This includes ancient obligate associations where the symbionts provide insects with essential nutrients and are strictly vertically transmitted through the host matriline. More widespread are heritable facultative symbionts, which are not essential for host survival but can provide important benefits such as expanding their hosts’ diet breadth, or conferring resistance to natural enemies, heat stress, and even pesticides [1–3]. Facultative symbionts are horizontally transferred between host lineages [4,5] and are often strongly non-randomly associated with particular host species or populations [6–9]. This raises the question of what factors determine the distribution of facultative symbionts across host species. One hypothesis is that insects tend to carry facultative symbionts that provide them with benefits that are specific to their ecological niche.

A common benefit provided by facultative symbionts is protection against natural enemies, such as nematodes, pathogenic fungi, viruses, and parasitoids [10–14]. It has been hypothesized that pressures from natural enemies may shape the distribution of protective symbionts across host species. If this were true, it would suggest facultative symbionts function in analogous manner to a horizontal gene pool from which insects can sample to rapidly adapt to changing pressures from natural enemies. However, physiological barriers to the horizontal transmission of facultative symbionts have also been identified [15,16], which may limit their spread.

One of the most extensively studied models for defensive symbiosis is the association between aphids and the facultative symbiont *Hamiltonella defensa*. *Hamiltonella* is known for being able to protect aphids from parasitoid wasp attack using toxins that are encoded on a phage that is integrated into the symbiont’s genome [4,17,18]. However, in certain aphid species, *Hamiltonella* can also be costly by decreasing host lifespan and increasing mortality [19,20]. Furthermore, it was shown that *Hamiltonella* genotypes differentially protect against different parasitoid species attacking aphids [15,21]. This opens the possibility that *Hamiltonella* distributions across aphid species may at least in part be shaped by parasitoids: aphids might host the *Hamiltonella* genotypes that are most efficient at protecting them against their most common natural enemies. However, the extent to which different selective pressures contribute to the genetic structure of *Hamiltonella* across host species is currently unknown.

To address this question, we genotyped the *Hamiltonella* strains in 412 aphids from 26 species using 6 housekeeping genes. Our survey revealed that *Hamiltonella* genotypes are not randomly distributed across aphid species, but rather form strong associations with particular hosts. To better understand the mechanisms explaining these associations we experimentally manipulated the infections status of 3 aphid species – *Acyrthosiphon pisum, Macrosiphoniella artemisiae* and *Macrosiphum euphorbiae* – using reciprocal transfection. Using a combination of fitness experiments, assessments of host-symbiont stability and parasitoid challenges, we tested whether *Hamiltonella* strains are: i) locally adapted to particular aphid species, and ii) whether aphids carry symbiont strains that provide strong protection against the parasitoid species most commonly attacking them.

## Results

### *Hamiltonella-aphid* associations are not random

We found that *Hamiltonella* genotypes are strongly non-randomly distributed among aphid species (Figure 1). We analysed the importance of *Hamiltonella* genotype, aphid species, aphid and *Hamiltonella* phylogeny, and the interactions between these variables, in explaining this pattern (Table S1). The only factor explaining the presence of *Hamiltonella* across host species was the host and symbiont co-phylogeny (BPMM: Posterior mode = 0.56, credible interval = 0.16 to 0.92, Table S1). This means that related aphid species tend to harbour a small number of related *Hamiltonella* genotypes. An example of this can be seen in the genus *Macrosiphum* where all species sampled were associated with a single clade of *Hamiltonella*, with one dominant genotype. The 187 aphids belonging to *A. pisum* (in Figure 1) were removed from this analysis due to this species being a genetically differentiated complex of plant-adapted biotypes that are known to harbour distinct *Hamiltonella* genotypes [7].

**Figure 1.**
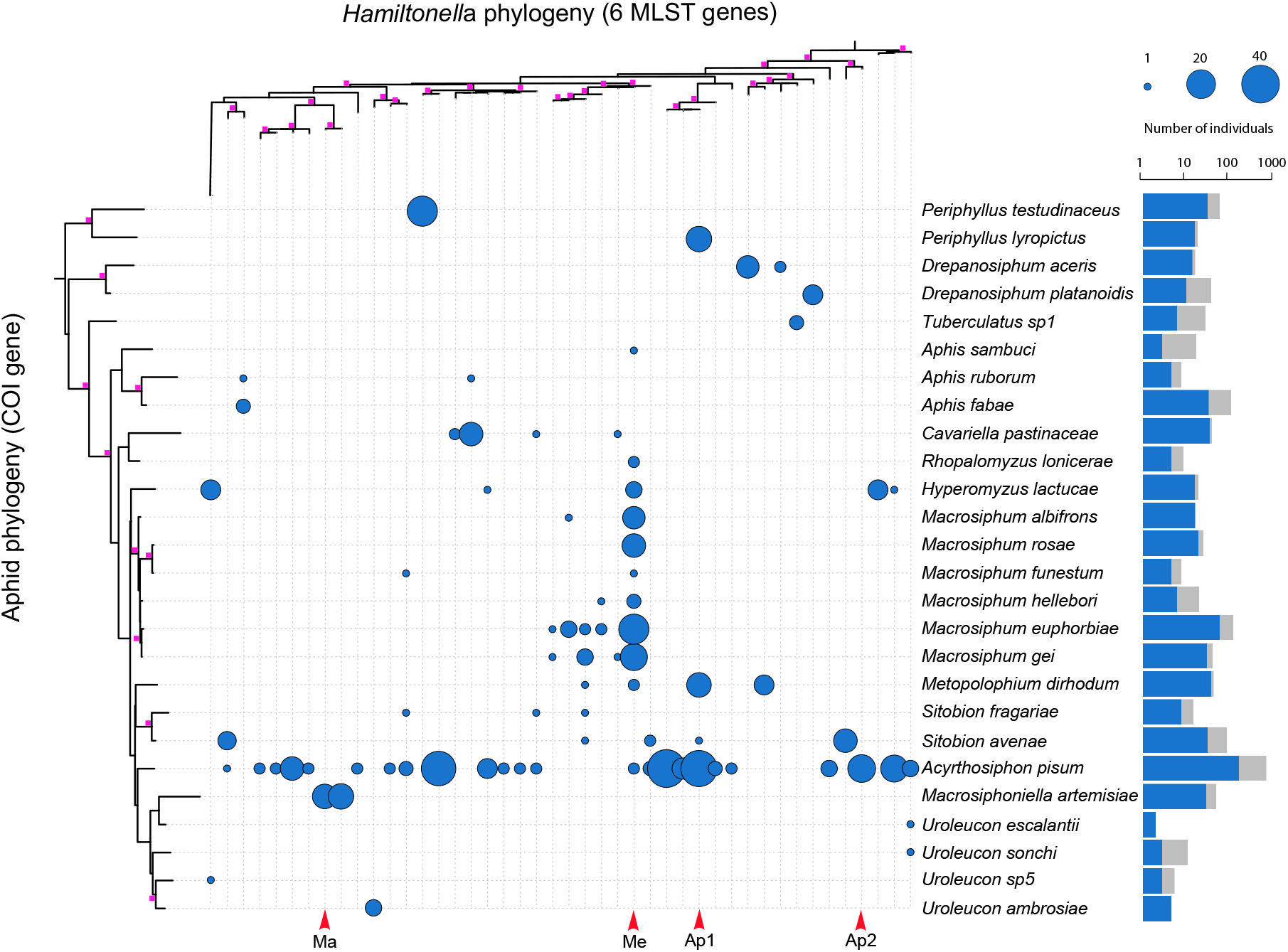
**Interaction matrix** of *Hamiltonella* genotypes (top phylogeny) occurring in aphid species (left phylogeny). Squares on the phylogeny nodes denote branch support >50. Bubble size corresponds to the number of times an aphid species was found harbouring a particular *Hamiltonella* genotypes. Barplot depicts the log_10_ total number of individual aphids screened (full bar) and the proportion carrying *Hamiltonella* (inset blue bar). Arrows at the bottom of the matrix denote *Hamiltonella* lineages used in the reciprocal transfection experiments.

### Mechanisms explaining *Hamiltonella-aphid* associations

To study the factors shaping host-symbiont associations, we focused on three aphid species (two clones each) that maintain strong relationships with specific *Hamiltonella* genotypes: *Macrosiphoniella artemisiae, Macrosiphum euphorbiae*, and the *Medicago* biotype of *A. pisum*. We studied four *Hamiltonella* strain: Ma is found exclusively in *M. artemisiae*; Me is the dominant strain associated with *M. euphorbiae* and all other *Macrosiphum* species surveyed; and Ap1 and Ap2 from *A. pisum*, one from each of the two major clades of *Hamiltonella* associated with the *Medicago* biotype [7] (Figure 1). To determine why each aphid species tends to harbour the *Hamiltonella* genotype(s) it does, rather than one of the genotypes found in the other species, we experimentally established aphid clones of each species carrying, its native *Hamiltonella* genotype(s), and those from other aphid species (i.e. non-native genotypes). We then assessed the impact of each *Hamiltonella* genotype-aphid species combination on i) the frequency the symbiont is passed to offspring, i.e. stability of maternal transmission, ii) the fecundity of aphids, and iii) protection against the parasitoid most commonly attacking the aphid species in nature. The reciprocal crosses resulted in 30 aphid *clone-Hamiltonella* genotype treatments: 3 aphid species * 2 clones * 5 infection status (4 *Hamiltonella* strains + 1 *Hamiltonella*-negative line) (Figure 2). We found very little difference between clones of the same species so results from both clones are presented and discussed together (results on individual clones can be found in the supplementary material).

**Figure 2.**
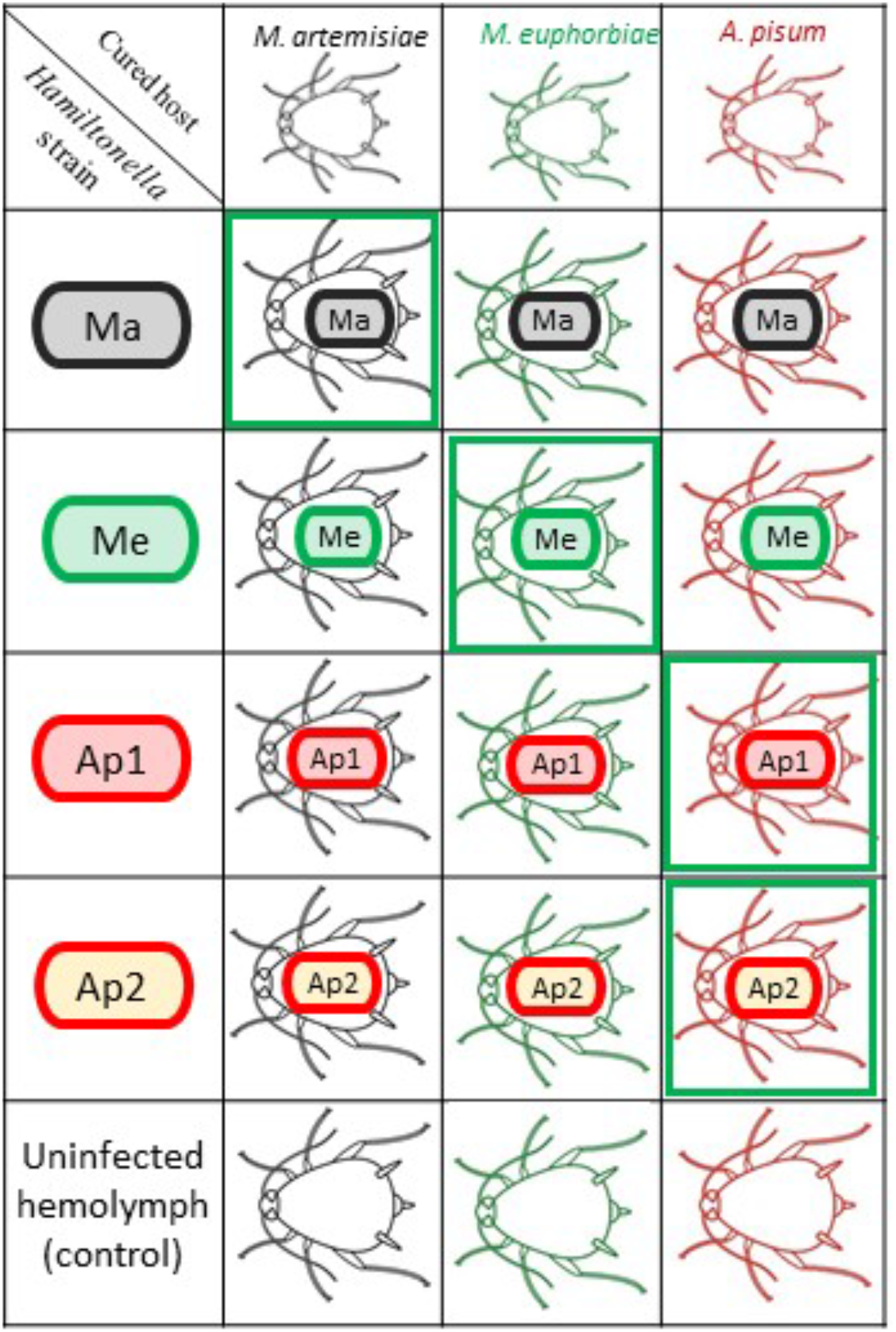
Cross-infection experimental design. We cured 2 clones each from 3 aphid species (*Macrosiphoniella artemisiae, Macrosiphum euphorbiae*, and the *Medicago* biotype - of *A. pisum*). Each clone was then reinfected with the symbiont genotype(s) it normally hosts in nature, i.e. its native genotype(s) (bold green boxes), and one of three non-native genotypes, or left cured as a control.

### *Hamiltonella-aphid* associations can be unstable and lethal to hosts

First, we assessed the stability of each experimentally established *Hamiltonella*-aphid combination by determining how often the symbiont is maternally transmitted to offspring. To do this, we tested for the presence of *Hamiltonella* in 3 offspring per newly infected female line for 5 generations using diagnostic PCR. Out of the 12 (24 including both clones) *Hamiltonella*-aphid species associations, 3 were found to be unstable, in that the symbiont was not perfectly transmitted to offspring (Figure 3, Table S2). In these cases, *Hamiltonella* was often rapidly lost from the clonal line: by generation 5, 100% (16/16) of *M. euphorbiae* lines injected with the *Hamiltonella* strain Ap2 had been lost. Similarly, *Hamiltonella* strain Ma was lost in both *A. pisum* (10/21 = 48%) and *M. euphorbiae* (6/13 = 46%) by generation 5. In all other combinations, no *Hamiltonella* loss was observed.

**Figure 3.**
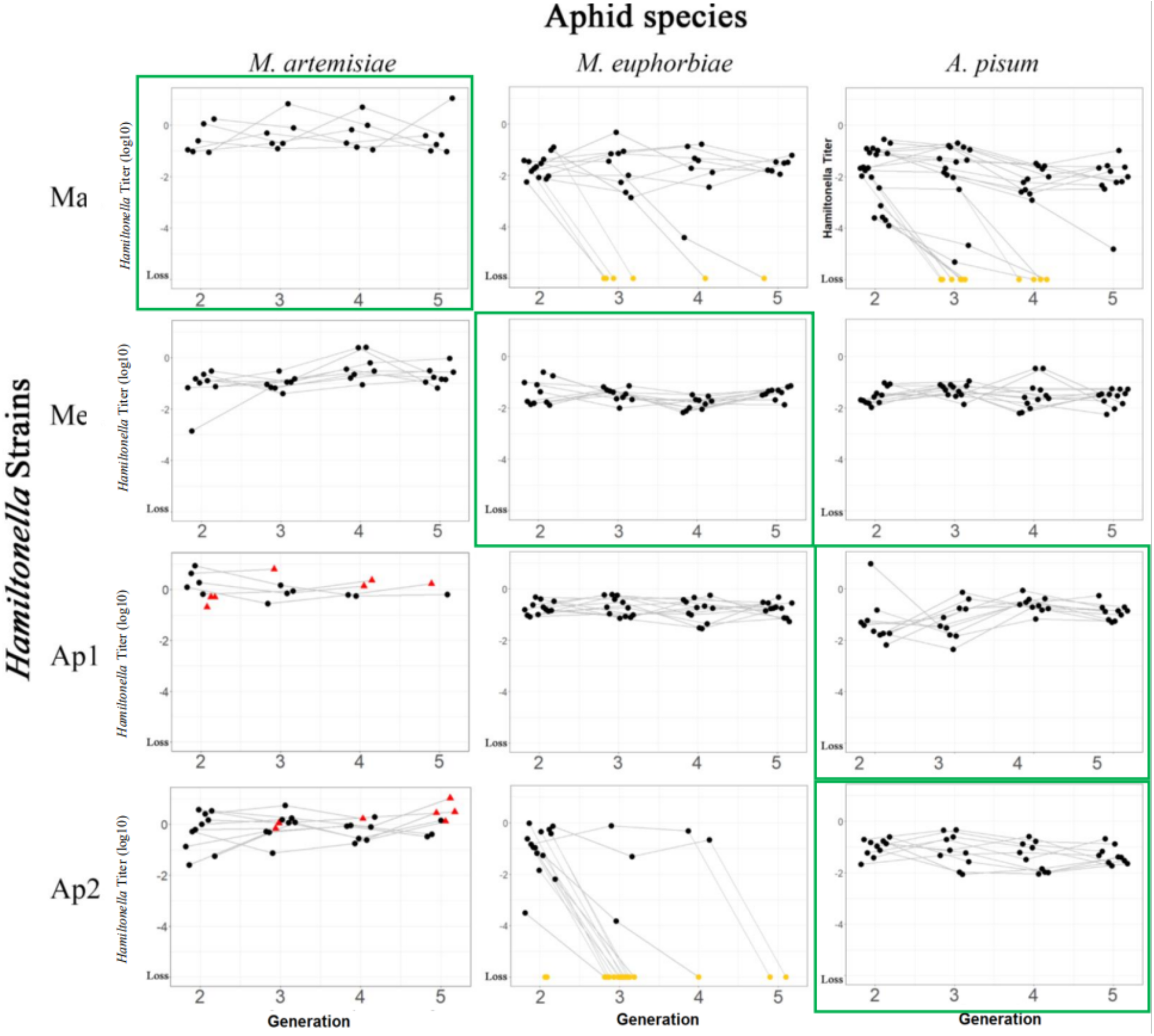
Changes in *Hamiltonella* titres across host generations following injection. Dots represent a single offspring (1^st^ of 3) screened from independent aphid lines injected with *Hamiltonella* with grey lines connecting the same aphid lineage across generations. Red triangles represent lineages that went extinct. Yellow dots at the bottom of the panels indicate the screened aphid no longer contained the symbionts, i.e. the symbiosis was lost. Bold green squares around panels highlight native host-symbiont combinations. Aphid lines that died in the same generation they were injected are not shown here. For the full data on symbiont losses in all offspring tested, line extinctions and initial establishment of infections, see Table S2.

In two additional associations, *Hamiltonella* strains Ap1 and Ap2 injected into *M. artemisiae*, symbionts were not lost, but rather led to early host death and a high rate of line extinction (triangles in Figure 3; Table S2): within 2 generations after the injection, 76% (16/21) of Ma clones infected with Ap1, and 50% (8/16) of those infected with Ap2, died. All Ap1-*M. artemisiae* and Ap2-*M. artemisiae* lines were extinct by generation 6 and 8, respectively. All unstable associations, and those that cause increased host mortality, were from aphid-*Hamiltonella* association not normally found in nature (i.e. non-native associations) (Figure 3, Table S2).

### Infection status impacts fecundity

We then assessed the impact of infection status on aphid fecundity (Figure 4). All fecundity assays were only conducted on a single clone per aphid species (Mug3 in *M. artemisiae*, PotG in *M. euphorbiae*, ApY in *A. psium*). The different *Hamiltonella* strains had no impact on the fecundity of *A. pisum* (χ^2^_4_=2.78, p=0.595) and *M. euphorbiae* (LRstat_4_=4.68, p=0.322).

**Figure 4.**
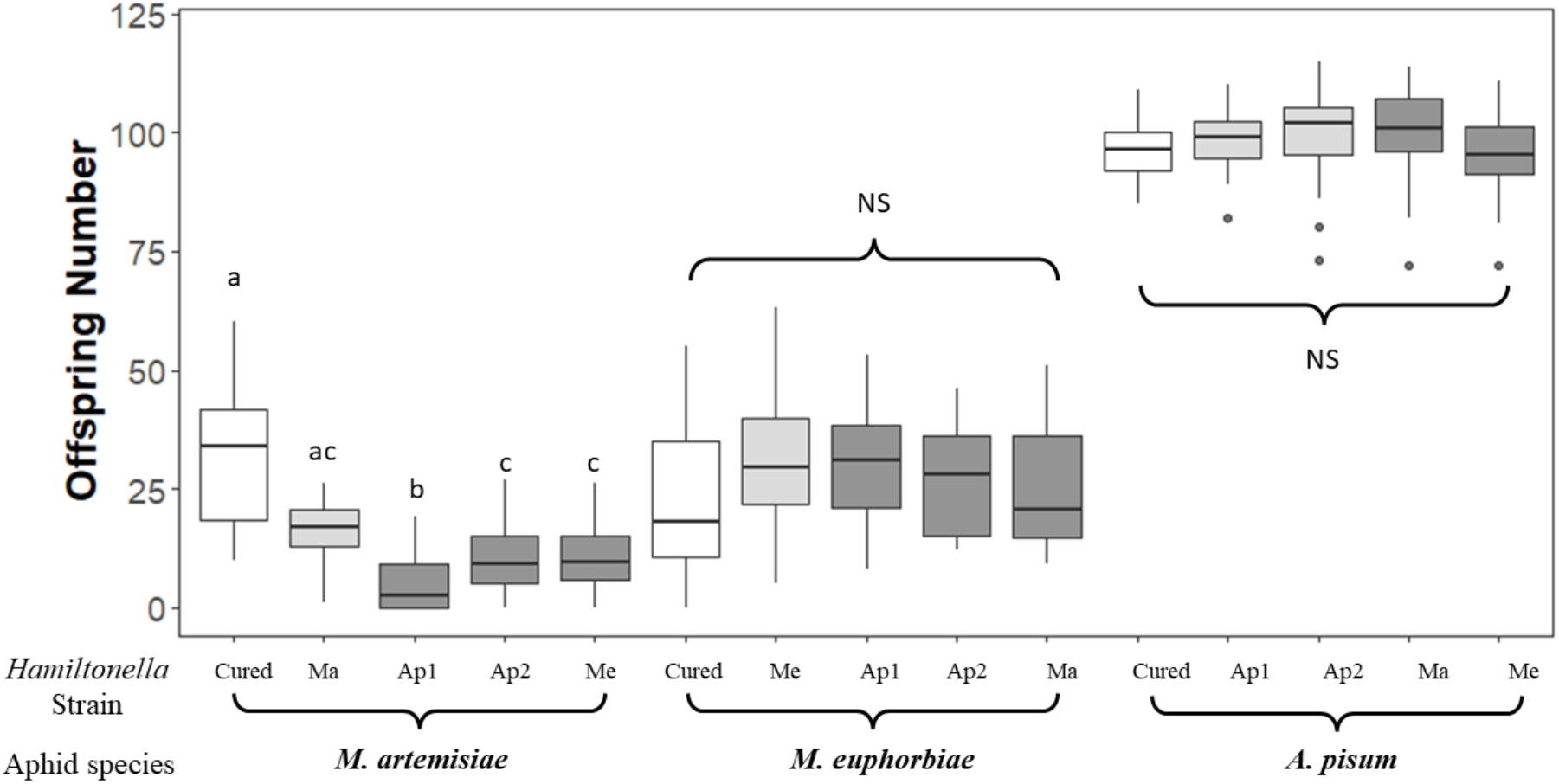
Impact of infection status on fecundity of *M. artemisiae, M. euphorbiae* and *A. pisum*. *Hamiltonella-negative* lines are shown in white, native associations in light grey, and non-native associations in dark grey. Fecundity assays were only conducted on a single aphid clone per species.

However, infection status had a strong impact on the fecundity of *M. artemisiae* (LRstat_4_=47.54, p<0.001). Interestingly, clones from *M. artemisiae* had the highest mean fecundity when uninfected with *Hamiltonella* (cured treatment; Figure 4). The native strain Ma did not significantly reduced *M. artemisiae’*s fecundity (z=-2.4, p=0.105), although Me did (z=-4.5, p<0.001). *Hamiltonella* strains Ap1 (z=7.2, p<0.001) and Ap2 (z=4.3, p<0.001) were the most deleterious to *M. artemisiae’* s fecundity, resulting in few offspring produced and extinction of the line (as shown in Figure 3).

### Infection status impacts resistance against parasitoids

Finally, we tested whether aphids are associated with *Hamiltonella* strains in nature that provide high degrees of protection against the parasitoid species that most commonly attack them. To test this hypothesis, we first surveyed the parasitoid species attacking the 3 aphid species used in this study. Second, we assessed the capacity of each *Hamiltonella* strains to protect aphids against the parasitoid species most commonly attacking them in the UK.

A total of 238 aphid-parasitoid associations were identified (Table 1 and Tables S3-S5) using a combination of deep-coverage barcoding of aphid mummies and sanger sequencing of parasitoids that emerged from morphologically identified aphids. All three aphid species in our study were predominantly attacked by a single parasitoid species: *Aphidius absinthii* in *M. artemisiae* (100%), *Aphidius ervi* in *A. pisum* (89%), and *Aphidius rhopalosiphi* in *M. euphorbiae* (67%). One parasitoid species, *A. ervi*, was found to attack both *A. pisum* and *M. euphorbiae* at lower frequencies (11%).

**Table 1.**
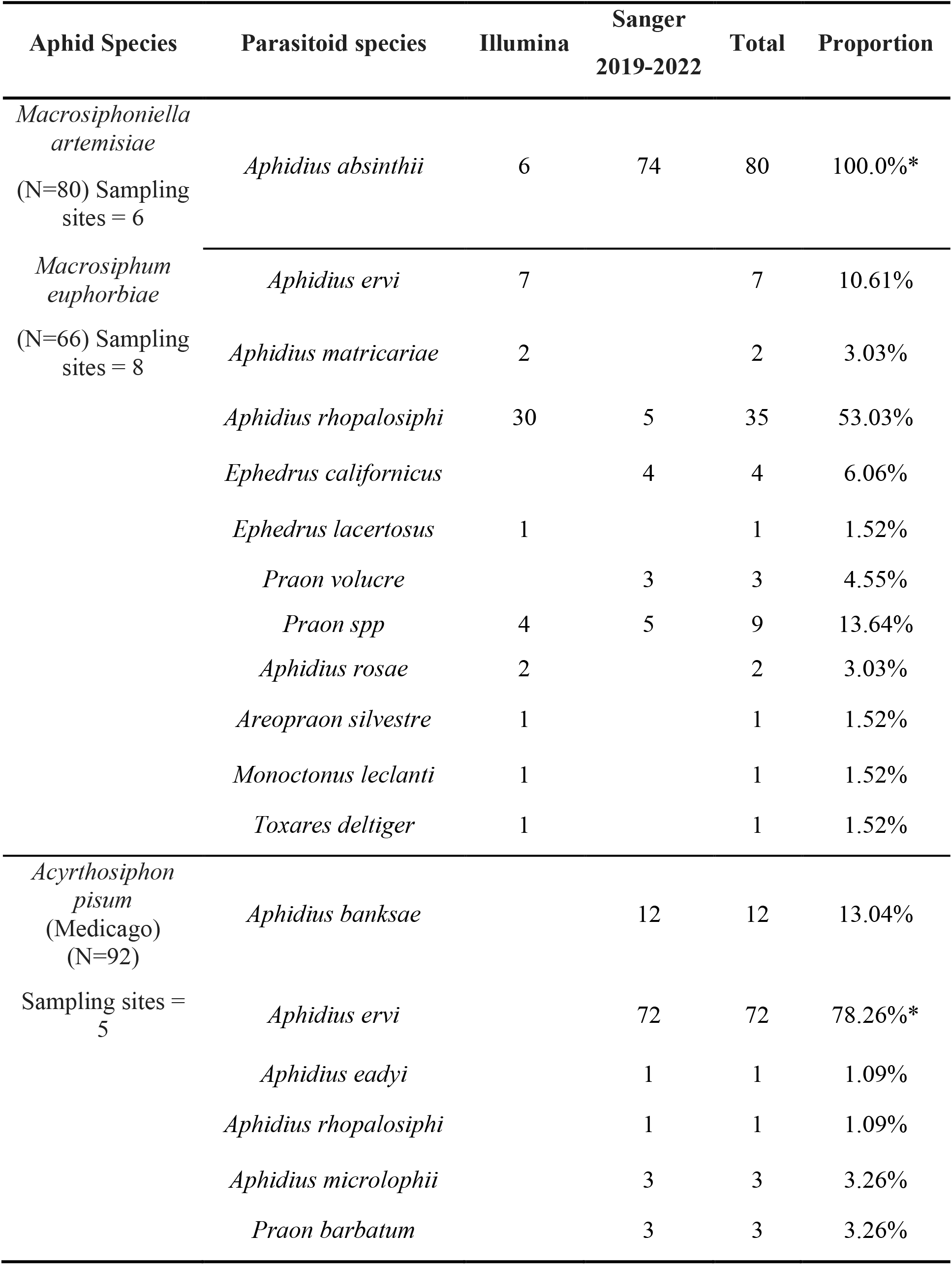
Aphid-parasitoid association list. Illumina sequencing was used for simultaneous identification of the aphid and parasitoid. Sanger sequencing was used to identify parasitoids emerging from aphids identified using morphology and host plant. Asterisks in the proportion column indicate the dominant parasitoid of each aphid. More information on collection can be found in Tables S3-5.

**Table 1.**
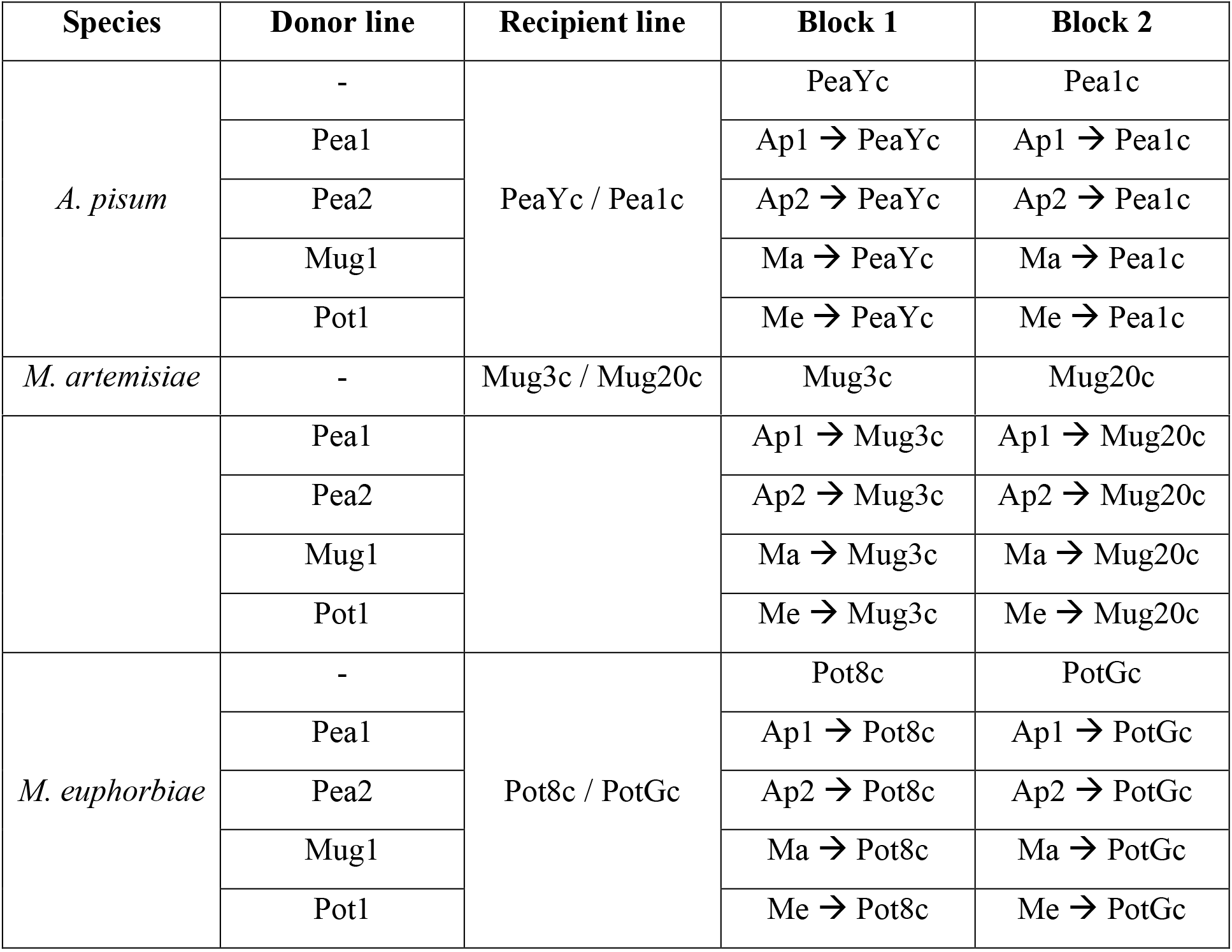
Aphid-*Hamiltonella* treatment lines obtained through curing and transfection.

Finally, we experimentally tested the impact of *Hamiltonella* infections status on resistance against parasitoids, by exposing each aphid species to the dominant parasitoid species attacking it. Infection status had a significant effect on mummification rate in all three aphid-parasitoid pairs: *M. artemisiae-A. absinthii* (χ^2^_2_=175.6, p<0.001), *M. euphorbiae-A. rhopalosiphi* (χ^2^_3_=641.5, p<0.001) and *A. pisum-A. ervi* (χ^2^_4_=1198.7, p<0.001). All cured aphids were highly vulnerable to attack by their dominant parasitoid, as shown by mummification rates close to 1 (Figure 5, Figure S1 for individual clones). When comparing the protection conferred by different symbiont strains, we found that in three of four cases, the native *Hamiltonella* genotype(s) provided the greatest degree of protection against the aphids’ dominant parasitoid: compared to other symbiont strains, Ma provided *M. artemisiae* with the highest degree of protection against *A. absinthii* (z=8.7, p<0.001), and both Ap1 and Ap2 protected *A. pisum* against its common parasitoid *A. ervi* (Ap1: z=-11.8, p<0.001, Ap2: z=-2.9, p=0.032), although one strain (Ap1) provided significantly greater protection (z=-13.7, p<0.001). In contrast, Me, the *Hamiltonella* strain most commonly found in *M. euphorbiae*, did not provide its host with any protection against *A. rhopalosiphi*. In *M. euphorbiae*, only the *A. pisum* derived Ap1 (z=-13.1, p<0.001) provided protection against the parasitoid *A. rhopalosiphi*. In fact, the Me *Hamiltonella* genotype did not provide protection against any parasitoid in any of the host backgrounds.

**Figure 5.**
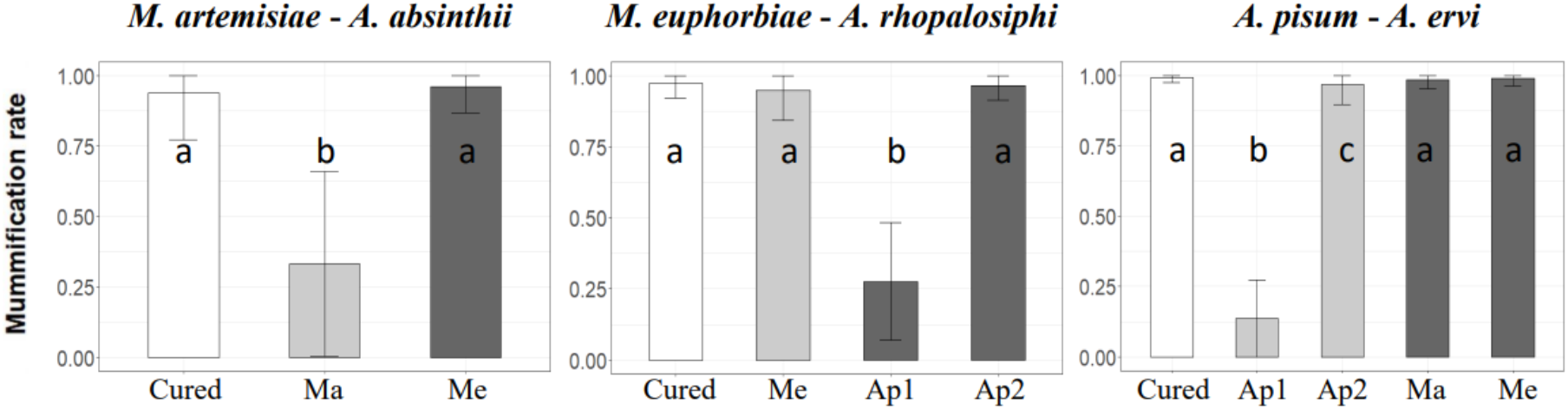
Effect of *Hamiltonella* genotype on protection from parasitoid wasp attack in three aphid species. Each aphid species was exposed to its dominant parasitoid. Protection was determined by the wasp mummification rate: 0 indicates that all aphids resisted the attack, and 1 that all aphids were parasitized (mean ± standard error). *Hamiltonella*-negative lines are shown in white, native associations in light grey, and non-native associations in dark grey. Only stable experimental lines were assessed. For results on individual clones see Figure S1.

## Discussion

### Explaining the non-random genetic structure of *Hamiltonella*

Our survey revealed that aphid species tend to form strong relationships with a single, or a few closely related, *Hamiltonella* strain(s). This is similar to what has been observed in pea aphid biotypes, where the plant-adapted aphids feeding on *Lotus*, *Medicago* and *Ononis*/*Melilotus* carry specific strains of *Hamiltonella* [7]. We also find intermittent cases where the same *Hamiltonella* genotype is carried by unrelated host species, which support previous finding that the symbiont is occasionally horizontally transferred between species [4]. This suggests that a combination of horizontal transfer and selection explain the observed patterns of *Hamiltonella* infections across aphid species. It has proposed that the facultative symbiont distributions of aphids may be the by-product of selection in response to attack by natural enemies [14], seasonal changes [22], and costs-benefits trade-offs [15,23,24]. However, the mechanisms explaining the strong genetic structure of *Hamiltonella* strains across aphid species have not been explored until now.

We found that in nearly all aphid-*Hamiltonella* combinations not normally found in nature, the symbiont strain was poorly adapted to either the aphid species or to the most common parasitoid species attacking the aphid. More specifically, we identified three selective filters that play a role in shaping the natural patterns of *Hamiltonella* infections across host species: i) symbiosis instability, ii) host lethality, and iii) lack of protective effect against the most common parasitoid of the host (summarized in Figure 6). In the case of the *A. pisum*-Ma association, both instability and lack of protective effect apply. By contrast, in native associations, the symbionts were always both efficiently vertically transmitted, avirulent, and in most cases, offered high degrees of protection against a prevailing nature enemy. This demonstrates that the *Hamiltonella* genotype associations found in nature tend to be locally adapted to both internal and external factors, i.e., to the aphid (genotype-genotype interaction) and to the aphid’s ecology (parasitoid pressure).

**Figure 6.**
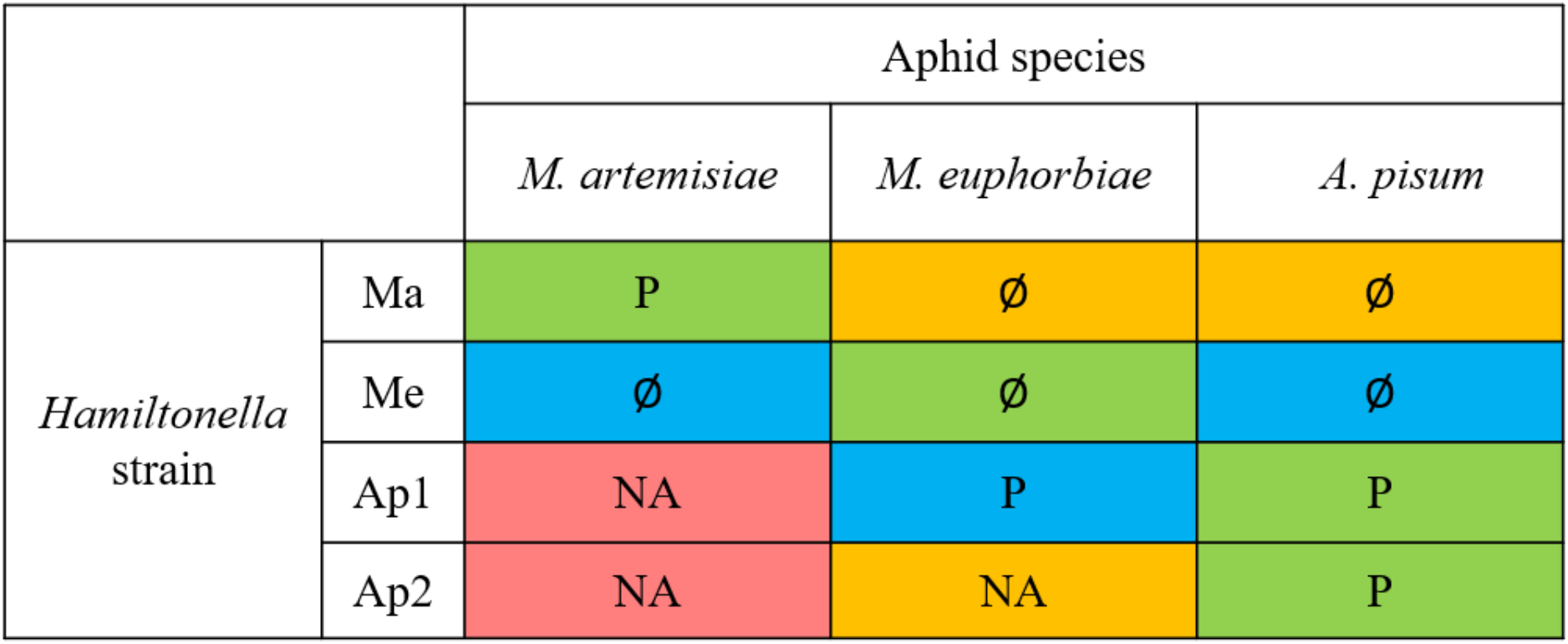
Summary of *aphid-Hamiltonella* experimental crosses. Native associations are shown in green background. Non-native associations are shown in blue unless they are unstable (yellow) or lethal (red). “P” indicates protection against the most common parasitoid, “Ø” indicates the absence of protection, and “NA” indicates that the level of protection was not assessed, due to the experimental association being unstable or lethal.

### Adaptation to internal factors

We assessed the compatibility of host-symbiont combinations by testing how reliably they transmit *Hamiltonella* to their offspring, and the fitness effects of different symbiont strains. Instability and lethality phenotypes were relatively consistent within clones of the same species, which suggests *Hamiltonella* genotypes were incompatible with certain host species (Table S2).

Studies on pea aphids have shown that facultative symbionts are more frequently lost during maternal transmission when transferred between host biotypes [25,26]. The probability of losses has also been shown to be influenced by the relatedness of host species [27–29]. In contrast, we found that all native host-symbiont combinations were perfectly maternally transmitted under laboratory conditions. Similar results have been cited for pea aphids, however, in other aphid species, such as in *Aphis craccivora*, maternal transmission of *Hamiltonella* is high but imperfect [30–32]. Imperfect transmission of facultative symbiont may also be exacerbated during sexual reproduction [33], and impacted by abiotic changes, such as temperature fluctuations [34].

Carrying facultative symbiont can also be costly in some host species, as a result of consuming a portion of the host’s resources [20,24,35]. However, cases of very strong virulence such as the ones we observed (Ap1 and Ap2 in *M. artemisiae*) have rarely been reported. Our findings are consistent with a previous study showing that *Hamiltonella* can be highly deleterious in *Aphis fabae*, with a decrease in fecundity up to 80% in certain host symbiont genotype combinations [36]. As with previous studies, we find that *A. pisum* can harbour diverse *Hamiltonella* strains with few costs to fecundity [37,38], and *M. euphorbiae* followed a similar trend [39], possibly even exhibiting a marginal, yet not significant, increase in fecundity when carrying *Hamiltonella*. Although previously thought to be largely benign, our results support the growing evidence that *Hamiltonella* infections can exhibit a wide range of host phenotypes - from avirulent to strongly virulent - with the outcome being determined by genotype-genotype interactions. Taken together, our results suggest the host-symbiont associations found in aphids are the product of co-evolution for decreased symbiont virulence and increased stability of maternal transmission. Furthermore, our results indicate that genetic incompatibilities are a major factor dictating the retention of symbiont genotypes across aphid species.

### Adaptation to external factors

Parasitoids are common natural enemies of aphids, and therefore a strong selective pressure for the evolution of resistance. Previous studies have shown that *Hamiltonella* strains provide specific protection against two distantly related species of parasitoids, *Aphidius ervi* and *Aphelinus abdominalis* [21]. Given that parasitoids tend to be highly hosts specific [40], we hypothesized that they may be a major force shaping protective symbiont genotype distributions across aphid species. We found that aphids tend to harbour *Hamiltonella* strains that provide them with at least some, to strong, levels of protection against the most common parasitoid species attacking them [1,41]. This supports our hypothesis that aphids tend to be associated with symbiont strains that provide them with high degrees of protection. Furthermore, we found that *Hamiltonella* strains can provide highly specific protection even against closely related parasitoid species. For example, the Ma strain of *Hamiltonella* provided protects against *A. absinthii*, but not *A. ervi*, which are both *Aphidius* parasitoids.

Our results suggest an aphids’ infection status is determined by the balance between the cost of carrying a *Hamiltonella* strain and the benefit resulting from protection. This theory suggests that horizontal transfer of symbionts is frequent enough that selection acts on aphids to retain highly protective symbiont stains. Alternately, selection may act on the symbiont itself to improve their protective effect in a host they are already adapted to.

In some cases, different aphid species share the same *Hamiltonella* genotype. This is most prominent in the *Macrosiphum* genus, but can also happen in more distantly related species, such as *A. pisum* and *Periphyllus lyropictus*. It would be interesting to know whether these species also share the same parasitoid species as this may explain the sharing of protective symbionts, through parasitoid-mediated horizontal transfer of *Hamiltonella*, and/or *Hamiltonella*-conferred protection against a shared natural enemy.

Our results suggest that pressures from natural enemies are an important force shaping the distribution of defensive symbiont genotypes found in nature. Future studies could test whether pressures from natural enemies are responsible for the genetic structuring of defensive symbionts in other systems. For example, in pea aphids, the *Hamiltonella* strains occurring in different biotypes may be the product of selection from different parasitoid species, or perhaps populations within a species, that are adapted to attacking the different aphid biotypes.

### Puzzling associations

For the most part, our results provide mechanisms that help explain why some aphid-*Hamiltonella* associations are found in nature and not others. However, the association of *M. euphorbiae* with Me, rather than Ap1, remains puzzling. Both strains are apparently well suited to the aphid, as they are both stable and avirulent, but contrary to Ap1, Me is not able to protect *M. euphorbiae* against *A. rhopalosiphi*, which is a common parasitoid attacking the aphid (a similar result was reported in *Aphis craccivora* [42]). As noted above, vertical transmission rates may differ in nature versus laboratory conditions, and this may help explain why Ap1 is not hosted by *M. euphorbiae*, if Ap1 is not effectively transmitted to offspring in nature. Alternatively, its absence in *M. euphorbiae* could be explained by the lack of opportunities for horizontal transfers to this host, or it may be an inferior intra-host competitor to the Me strain. However, the prevalence of Me in *M. euphorbiae* (and other *Macrosiphum* species) at such high frequencies (Fig. 1) is in itself surprising, as we did not find evidence for any benefit conferred to its host.

*Macrosiphum euphorbiae* tends to be attacked by a greater diversity of parasitoid species and to be associated with more *Hamiltonella* strains than the other aphid species. The lack of protection conferred by Me is therefore arguably less surprising than in a species consistently attacked by only one parasitoid species (such as *M. artemisiae*). First, based on collections mostly performed on *Geum* spp. and *Gallium aparine* (Table S4), we found that *M. euphorbiae* was most commonly attacked by *A. rhopalosiphi*. However, it is possible that on other plants, *M. euphorbiae* is attacked by different parasitoids that Me protects against. The rate at which aphids are targeted by a given parasitoid has been shown to vary depending on the host plant [43] For example, *M. euphorbiae* feeding on pepper plants in Spain have are more commonly attacked by *Aphidius colemani*, *Praon volucre* and *Aphidius matricariae* [44], whereas in North America potato farms, it is primarily attacked by *Aphidius nigripus* [45]. Second, *M. euphorbiae* can harbour several related *Hamiltonella* strains, and we only tested one of them (the most common, according to our survey). It is possible that other strains provide protection against *A. rhopalosiphi*. As a result, we cannot exclude the possibility that the *Hamiltonella* association found in *M. euphorbiae* is, to some extent, influenced by symbiont-parasitoid associations that we did not test in our study.

The *M. euphorbiae*-Me association might also be explained by adaptation to external factors unrelated to parasitoids. It might, for example, protect against non-parasitoid natural enemies, e.g. lady beetles, as reported by [46], or suppress the immune system of host plants [47]. It is therefore possible that although Me did not seem to improve *M. euphorbiae* performance on broad beans, it may increase its fecundity on other host plant species, including its native host plant *Gallium*.

### The horizontal gene pool hypothesis

It has been suggested that secondary symbionts function as a horizontal gene pool that insects can draw from to rapidly adapt to changing environments [7]. Our results suggest that the access to this horizontal gene pool is restricted by barriers to the establishment of new associations, in the form of genotype-genotype interactions (see also [20]). However, the occurrence of the same *Hamiltonella* genotypes at low frequencies in different hosts does support on-going horizontal transfer among aphid species. Our results demonstrate that in several cases, non-native strains can lead to stable, avirulent and beneficial association that could be retained (e.g. Ap1 in *M. euphorbiae*). Additionally, 2 of 3 ‘unstable’ combination in our study were not completely lost, only some of the infected lines lost *Hamiltonella*. Is it possible that associations such as these may stabilise after a period of coevolution, in particular if they provide a selective advantage, such as parasitoid protection, that retains them in a population? Furthermore, the potential for rapid adaptation is supported by our finding that the Ap1 strain is able to efficiently protect *M. euphorbiae* against *A. rhopalosiphi*. As parasitoid frequencies change over time, these genotypes, if protective, might allow for rapid adaptation and approach fixation. Taken together, our results provide strong support for the local adaptation of aphid species and specific *Hamiltonella* strains and suggest that changes in pressure from parasitoids may be rapidly addressed by the acquisition of new symbiont strains

## Material & Methods

### Patterns of *Hamiltonella*-aphid associations

#### Aphid collection and identification

Aphids were collected in the UK between 2011 and 2019 by beating plants over a white tray or manually removing them from plants, before being placed in 100% ethanol. Resampling of the same aphid clones was minimized by separating collections from the same plant species by at least 10 m. Aphids were identified by DNA barcoding based on data from [48] and confirmed using morphological examination following [49]. Genomic DNA was extracted from individual specimens using DNeasy Blood and Tissue kits (QIAGEN, Venlo, Netherlands) and then we amplified an approximately 700 bp DNA fragment of the cytochrome c oxidase I (COI) mitochondrial gene from the DNA using the Lep F and Lep R primers. We sequenced the amplicons in the forward direction (Full details of PCR conditions and primer sequences are provided in Table S6). DNA sequences were aligned with MUSCLE (https://www.ebi.ac.uk/Tools/msa/muscle/). Aphids were identified to species by comparing COI sequence data to the online databases BOLD (http://www.boldsystems.org/) and GenBank using BLAST.

#### Hamiltonella *screening and genotyping*

*Hamiltonella* was detected using diagnostic PCR based on the 16S ribosomal RNA gene (Table S6) amplified from the whole body aphid DNA extracts. Genotyping was performed using a multilocus sequence-typing (MLST) scheme containing six bacterial housekeeping genes: accD, gyrB, hrpA, murE, recJ, and rpoS [50]. PCR products were Sanger-sequenced in the forward direction, and aligned using MUSCLE (https://www.ebi.ac.uk/Tools/msa/muscle/).

#### Phylogenetic reconstruction

Host (COI) and symbiont (MLST) phylogenies were reconstructed using PhyML 3.0 (http://www.atgc-montpellier.fr/phyml/) with 100 standard bootstrap analysis and otherwise default parameter values. *Adelges cooleyi* and a *Hamiltonella* strain of *Bemisia tabaci* were used as outgroups in the host phylogeny and the symbiont phylogeny, respectively.

#### Bayesian general linear modelling

We used Bayesian general linear models (BPMMs) with Markov Chain Monte Carlo (MCMC) estimation run in the mixed model package MCMCglmm [51] in R v4.2.0 [52].. The occurrence and non-occurrence of *Hamiltonella*-aphid combinations were fitted as a binomial response variable. *Hamiltonella* genotype, aphid species, genotype-species interaction, *Hamiltonella* phylogeny, aphid phylogeny, and *Hamiltonella*-aphid cophylogeny were included as random explanatory variables.

The MCMC was run for ten million iterations with a thinning interval of 225 and a ‘burn in’ of 100000. Convergence of the chains was confirmed by visual inspection of the trace plots. We present the results as the posterior modes (PM) with the 95% credible intervals (CI) of the estimate.

### Experimental and maintenance of aphid lines

#### Establishment of aphid lines

Aphids belonging to three different species (*A. pisum*, *M. artemisiae*, & *M. euphorbiae*) were collected live in the field. Three clonal lines per species were established. Secondary symbionts infection statuses of all aphid clones were tested by 16s rRNA PCR (Table S6). They were found to be positive for *Hamiltonella*, but negative for *Serratia symbiotica*, *Fukatsuia symbiotica, Rickettsia* sp., *Rickettsiella* sp. and *Spiroplasma* sp. and *Regiella insecticola*. Aphids cured of *Hamiltonella* using selective antibiotics served as “recipients” for transfection. Clonal lines carrying each *Hamiltonella* strain of interest were retained as symbiont “donors” for artificial transfections (Table 1). Clonal lines of aphids were maintained in the lab at 15°C with a 16h light (Sylvania Gro-Lux F36W/GRO-T8 bulb) 8h dark cycle on a leaf of *Vicia faba* (*A. pisum* and *M. euphorbiae*) or *Artemisia vulgaris* (*M. artemisiae*) embedded in 2% agar in a Petri dish. Leaves were changed weekly.

#### Antibiotic curing

We used antibiotic treatments to selectively remove *Hamiltonella* without eliminating the primary symbiont *Buchnera aphidicola*, following a protocol adapted from [53]. The antibiotic solution was obtained by mixing 10 mg/mL of Ampicillin sodium salt, 5 mg/mL Cefotaxime sodium salt, and 5 mg/mL Gentamicin in water. A single leaf of the host plant was cut and placed in a 0.5 mL Eppendorf tube filled with the antibiotic solution. We placed 10 one- or two-day old aphid nymphs on a leaf and left them to feed for five days on antibiotic solution. Surviving aphids were then transferred to a regular Petri dish describe above. We confirmed the antibiotic had removed *Hamiltonella* by testing aphids in the second generation after treatments, and in the sixth generation prior to the start of experiments, using symbiont-specific 16S rRNA primers. The presence/absence/strain of each *Hamiltonella* treatment was reconfirmed after the experiments had been conducted.

#### Hamiltonella transfections

*Hamiltonella*-aphids associations were established using haemolymph injection (Figure 2, Table 1). Approximately 0.25 μl of haemolymph was obtained by removing the leg of a naturally infected adult (donor) aphid and then injected into a first instar uninfected (recipient) aphid using a microcapillary needle [15]. Injected aphids were maintained until they reach adulthood and the presence of *Hamiltonella* was checked in 3 of their late offspring (>10^th^ in birth order, in most cases) through DNA extraction and PCR (as described above). Successful injection lines were kept for a minimum of seven generations before being used in parasitoid experiments, to ensure the stability of the infection [54]. At least 13 independent injection lines per *Hamiltonella*-aphid combination were established (Table S2). Their infection status was confirmed both immediately before and immediately after all experiments in which they were involved using 16S rRNA PCR by testing the siblings of the aphids being assayed.

### Stability of *Hamiltonella* infections and fecundity effects

#### Stability of Hamiltonella infections

We assessed the stability of newly established *Hamiltonella* infections at generations 2 to 5 following injection by measured the relative density of *Hamiltonella* using quantitative PCR on whole aphid DNA extracts. Three 14-days-old aphids per injection line (6 injection lines per *Hamiltonella*-aphid combination) were extracted as described above. We used two single copy genes: one in the aphid nuclear genome (EF 1-alpha) and one in *Hamiltonella* (dnaK). The quantification was performed on a CFX Connect Real-Time PCR Detection System (BioRad, Hercules, California, U.S.A.). Full details of PCR conditions and primer sequences are provided in Table S6. The mean qPCR efficiencies were calculated using a ten-fold series of dilutions from 3.2 x 10^2^ to 3.2 x 10^7^ copies of purified PCR products. The efficiencies were 96.3% for the aphid gene and 89.1% for the *Hamiltonella* gene. Samples were run in triplicates. As the standard deviations between the triplicates of a given samples were below 0.5 cycles, the mean quantification cycle (Cq) values were used to calculate the starting quantities of the genes of interest. For each sample, the starting quantity for the *Hamiltonella* gene was divided by the starting quantity for the aphid gene to obtain the *Hamiltonella* density.

#### Aphid fecundity

Lifetime fecundity was recorded for 5 to 36 (~24 on average) adult aphids per infection status and recipient species (Mug3, PotG and PeaY), from either cured lines or newly injected lines (from generation 2 to generation 7 following microinjection). Dishes were checked weekly for offspring. Aphids that died prematurely from fungal infection were excluded from the dataset.

### Parasitoid survey and resistance experiment

#### Collection and identification of mummified aphids

Mummified and live aphids were collected in the Greater London area (UK) on 5 plant genera or species (*Geum* spp., *Galium aparine, Medicago sativa* and *Artemisia* spp.) known to host *M. euphorbiae*, *A. pisum*, and *M. artemisae* (see Table S4 and S5 for full information on mummy collection). Mummified aphids were immediately preserved in 70% ethanol for DNA extraction. Live aphids were kept in the lab on a leaf of their host plant for two weeks. Any mummies forming during this time was either preserved for DNA extraction or used to establish lab colonies (see below). DNA extraction of mummified aphids was performed as previously described.

To simultaneously identified the aphid and its parasitoid from field collected aphids and mummies, we amplify DNA using universal barcoding primers that target the cytochrome c oxidase subunit I ‘COI’ gene, Ill_B_F/HCO2198 [55] (Table S6). Tagged PCR products were submitted to Bart’s and the London Genome Centre for addition of indices and pooling of libraries. Sequencing was then carried out on a single MiSeq run (paired-end, 2 x 300 bp reads). Samples were analysed using Dada 2 v1.16 [56].

#### Parasitoid resistance experiment

Colonies of three parasitoid species were maintained in the lab on host aphid clones that had been cured of *Hamiltonella: Aphidius absinthii* on *M. artemisiae* (Mug3c), *A. ervi* on *A. pisum* (PeaYc) and *A. rhopalosiphi* on *M. euphorbiae* (PotGc). Temperature and lighting were the same as for aphids. Prior to the experiment, newly hatched wasps of both sexes were kept together for at least 24 hours to allow for mating. Parasitoid females were then individually exposed to one second-instar larva from the tested aphid line. In case the parasitoid failed to attack the aphid within 10 minutes, it was discarded. Otherwise, it was transferred to a Petri dish containing 15 second instar aphids (4-day-old instars for *M. euphorbiae* and *A. pisum*; 6-day-old instars for *M. artemisiae*) and kept there for 24 hours. After 12 days, the mummification rate was calculated as the number of mummified individuals divided by the number of individuals that were either alive or mummified (non-mummified aphids that died before 12 days were excluded). At least 12 biological replicates were used for every experimental line tested.

### Statistical analyses

Statistical analyses were performed in R v4.2.0 [52] using RStudio v1.4.1743 [57]. The lme4 (v1.1-29) package [58] was used to fit generalised linear mixed models (GLMM). The multcomp (v1.4-19) package [59] was used to perform post-hoc tests.

Fecundity data were analysed independently for each species. Three GLMs were fitted, one (*A. pisum*) with a Poisson distribution, two (*M. artemisiae* and *M. euphorbiae*) with a negative binomial distribution (as overdispersion prevented the use of a Poisson distribution). Infection status was included in the model as a fixed effect, and injection line was nested within infection status. Post-hoc tests were performed after model selectioine numbers.

Mummification rate data were analysed independently for each species, by fitting three GLMMs with binomial distributions. Infection status was included as a fixed effect, and clone as a random effect. Post-hoc tests were performed after model selection.

## Data accessibility

Data and scripts are available online: https://doi.org/10.5281/zenodo.6784841. The GenBank accession numbers for the MLST, and COI gene sequences determined in this study are ###### to ######.

## Author’s contribution

T.W.: conceptualization, investigation, formal analysis, writing – original draft, writing – review and editing; D.M.: conceptualization, formal analysis, writing—original draft, writing—review and editing; R.A.L.: investigation, writing—review and editing; L.M.H: conceptualization, funding acquisition, investigation, supervision, writing – original draft, writing – review and editing.

## Acknowledgments

We would like to thank Susie Hawthorne and Timothy Penny for their help with experiments and analyses. We would also like to thank the members of the Henry lab for their feedback.

