## Supplementary material for "Local adaptation to hosts and parasitoids shape *Hamiltonella defensa* genotypes across aphid species": Figure S1

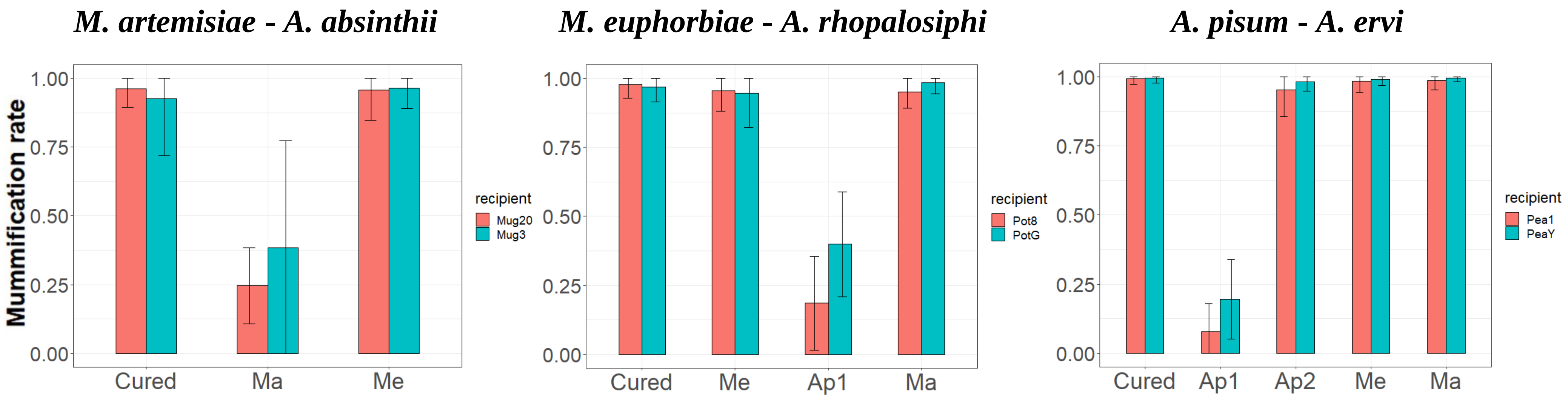


**Figure S1. Effect of *Hamiltonella* genotype on protection from parasitoid wasp attack in 6 aphid clones.** Two aphid clones from each aphid species were exposed to its dominant parasitoid. Protection was determined by the wasp mummification rate: 0 indicates that all aphids resisted the attack, and 1 that all aphids were parasitized (mean ± standard error). Only stable experimental lines were assessed. Two different clones per aphid species are coloured in red and blue.
